# The genome of the Glacier lanternfish shows loss of MHC I and II function and provides insight into evolution of lanternfish immune systems

**DOI:** 10.1101/2025.08.28.672920

**Authors:** Monica Hongrø Solbakken, Ole K. Tørresen, Ave Tooming-Klunderud, Morten Skage, Torkild Bakken, Kjetill Sigurd Jakobsen

## Abstract

Lanternfishes (Myctophidae) are one of the most abundant and species diverse orders inhabiting the mesopelagic zone. Exploitation of marine resources has recently attracted increased interest. It is therefore essential to improve our understanding of the meso- and deep pelagic ecosystems with respect to conservation and management strategies. Genomic resources are paramount to enable in-depth studies of population structuring and understanding the inherent genetic diversity, but also to enable studies of processes such as local adaptation and selection. Here we present a genome assembly of the Glacier lanternfish (*Benthosema glaciale)* generated with long-read PacBio and Hi-C contact map data. By comparing additional available genomes from the Myctophiformes order we explore their adaptive immune strategies. Our findings reveal that multiple lineages within this order have lost a significant proportion of genes related to adaptive immunity (*Electrona, Protomyctophum* and *Benthosema*). We find simultaneous loss of both classical MHC class I and class II function coinciding with reduction in genetic diversity with respect to immunoglobulin and T-cell receptors. In contrast, the sister branch represented by *Gymnoscopelus* and *Nannobrachium* appear to have maintained the standard configuration of the jawed vertebrate adaptive immune system apart from large gene expansions in MHC class II. Our results demonstrate that the *Benthosema*-belonging lineage (Myctophinae) has completely lost the core functions of the adaptive immune system. How, when and why this occurred warrants further investigations.

## Introduction

The mesopelagic zone contains a substantial amount of oceanic biomass and is gaining increasing attention with respect to potential exploitation [1]. Currently, the main usage of this resource is production of feed and oil for aquaculture [1], but there are efforts exploring future use in human consumption and nutrient supplements as well as pharmaceutical and healthcare related bioprospecting [2–5]. Furthermore, some species have been proposed suitable for studying and monitoring levels of anthropogenically derived contaminants such as microplastics [6, 7].

Mesopelagic fish are centrally located in the food web being both predator and prey. They also take part in the ocean carbon pump as many of the species perform diel vertical migration to forage in shallower waters [1, 8]. Given our limited understanding of the mesopelagic zone, the increased interest in this marine resource has prompted further intense research especially within conservation and management [1, 5]. However, there are large knowledge gaps including species diversity, spatial distribution, population structure, prey choices, trophic interactions and temperature adaptations. Currently, exploitation of this resource has been met with a precautionary tone given the potential impacts on the broader ocean ecosystem [1, 5, 9].

Myctophidae (Lanternfishes) and Gonostomatidae (Bristlemouth/Lightfish) dominate the mesopelagic zone with an overall biomass estimated between 1.8 and 15.9 Gt [1, 5, 10]. Myctophidae is part of the order Myctophiformes together with the Neoscopelidae (Blackchins). In total, the order contains around 250 species where the majority belongs to the highly diversified Myctophidae [11, 12]. Myctophidae are generally small fish that inhabit marine waters across the globe and reside in surface waters to the upper range of the bathypelagic zone (1000-4000m). Lanternfishes have a lifespan between 1 to 5 years, but show low fecundity rates (100-2000 eggs) [10]. The name lanternfish stems from their coelenterazine-based photophores which are arranged in species-specific patterns along their bodies and are believed to aid in species identification, communication and counter-illumination [12–14]. While photophores in Neoscopelidae are most likely used as counter-illumination and camouflage, the Myctophidae photophores have migrated to a more lateral position where patterns are species-specific [15, 16]. Several studies suggest that the photophores contribute to the formation of genetic isolation of populations or ecotypes given the exceptional species diversity within the family [15–17]. This also overlaps a great diversification with respect to photophore morphology [18], visual adaptations [19] mouth size [20], body size and otolith morphology [21, 22], feeding strategy [23], adaptations to oxygen minimum zones [10], as well as dentition [24]. Efforts to characterize the species diversity is further complicated by cryptic speciation and poorly resolved phylogenetic relationships both using morphological and genetic markers [12, 15, 16, 22, 25–29].

The Glacier lanternfish *Benthosema glaciale* (Reinhardt, 1837, *B. glaciale*) is widely distributed in the North Atlantic Ocean, particularly in the fjords along the Norwegian coast, in the Norwegian Sea north of Svalbard as well as in the Mediterranean Sea [30–39]. In Norway, *B. glaciale* is among the most abundant species in the deep fjords [4]. A study with limited genetic data indicates three distinct populations in the Atlantic Ocean [40]. Like many other Myctophidae, *B. glaciale* also performs diel vertical migration where feeding patterns and diet correlate with age, season, geographic range and overall size [30–32]. This species is currently being explored as an emerging resource for aquaculture feed and potentially human consumption [2–4]. However, improved genomic resources such as a high-quality reference genome effectively allowing population and functional studies will allow for a deeper insight into its ecology, evolution and general biology.

Here we present a high quality long-read genome assembly of *B. glaciale* as a common resource and tool for further investigations of this species and its relatives. In particular, we explore the comparative genomics of the adaptive immune system using the *B. glaciale* assembly and some additional available Myctophiformes genomes. Our findings show loss of classical Major Histocompatibility Complex (MHC) class I and class II function in *Benthosema, Protomyctophum* and *Electrona*, but not in *Nannobrachium and Gymnoscopelus*, highlighting that immune-genomic differences may be very important to consider in overall management strategies for this group of ecologically and economically important species.

## Results

The genome of *B. glaciale*, had an estimated genome size of 861 Mb, with 1.96% heterozygosity and a bimodal distribution based on the k-mer spectrum (**Supplementary Figure** 1). A total of 26-fold coverage in Pacific Biosciences single-molecule HiFi long reads and 98-fold coverage in Arima Hi-C reads resulted in two haplotype-separated assemblies. The final assemblies have total lengths of 1282 Mb and 1301 Mb, respectively. Pseudo-haplotypes one and two have scaffold N50 size of 45.0 Mb and 45.0 Mb, respectively, and contig N50 of 0.48 Mb and 0.47 Mb, respectively (**Figure 1**). 22 autosomes were identified in both pseudo-haplotypes (numbered by length in pseudo-haplotype one with the homolog in pseudo-haplotype two receiving the same number). In pseudo-haplotype two, chromosome 1 is split in two, termed 1A and 1B (**Supplementary table 2**). The BUSCO scores were 95.8% for pseudo-haplotype 1 and 96% for pseudo-haplotype 2 (**Figure 1**). 33,856 protein-coding genes were annotated in pseudo-haplotype one, and 34,138 pseudo-haplotype two, with 95.2% and 95.4% BUSCO completeness, respectively (**Supplementary table 2**).

The adaptive immune-genomic structure of lanternfishes (Myctophidae) was investigated by manual annotation of key genes related to antigen presentation (MHCs), antigen recognition (T-cell receptors, TcR) and antibody production (B-cell receptors, heavy and light chain immunoglobulins) in the *B. glaciale* genome together with seven publicly available Myctophidae genomes including *Benthosema pterotum, Electrona antarctica, Gymnoscopelus braueri, Gymnoscopelus microlampas, Nannobrachium achirus, Protomyctophum bolini* and *Protomyctophum parallelum* (**Table 1** and **Table 2**).

**Table 1.** Genomes downloaded and generated for this study.

| Scientific name (FishBase) | Common name | Subfamily (FishBase) | Distribution (FishBase) | Assembly name | Assembly NCBI accession | Assembly owner | Genome size | Chromosomes | Genome coverage |
| --- | --- | --- | --- | --- | --- | --- | --- | --- | --- |
| *Gymnoscopelus braueri* | Brauer's lanternfish | Gymnoscopelinae | Generally ranges between the coasts of Antarctic and 33°S - 46°S | fGymBra2.1 | GCA_963280865.1 | Wellcome Sanger Institute | 1.4 Gb | 24 | 34x |
| *Gymnoscopelus microlampas* | Minispotted lanternfish | Gymnoscopelinae | Circumglobal between Subtropical Convergence and Antarctic Polar Front | fGymMic1.1 | GCA_963454915.1 | Wellcome Sanger Institute | 1.3 Gb | 24 | 62x |
| *Nannobranchium achirus* (*Lampanyctus achirus*) | Cripplefin lanternfish | Lampanyctinae | Circumglobal from the Subtropical Convergence to south of the Antarctic Polar Front | fNanAch1.1 | GCA_963921795.1 | Wellcome Sanger Institute | 1.7 Gb | 24 | 31x |
| *Electrona antarctica* | Antarctic lanternfish | Myctophinae | Circumpolar south of Antarctic Polar Front | fEleAnt2.1 | GCA_951216825.1 [42] | Wellcome Sanger Institute | 1.4 Gb | 24 | 43x |
| *Protomyctophum bolini* | Bolin's lanternfish | Myctophinae | Circumpolar between Antarctic Divergence and Subtropical Convergence zone | fProBol1.1 | GCA_963924005.1 | Wellcome Sanger Institute | 1.1 Gb | 24 | 40x |
| *Protomyctophum parallellum* | Parallel lanternfish | Myctophinae | Circumglobal between the Subtropical Convergence and the Antarctic Polar Front | fProPar1.1 | GCA_964188405.1 | Wellcome Sanger Institute | 1.2 Gb | 23 | 35x |
| *Benthosema pterotum* | Skinnycheek lanternfish | Myctophinae | Indo-West Pacific, Southeast Atlantic | ASM3910535 v1 | GCA_039105355.1 [43] | Xiamen University | 1.3 Gb | 24 | 88x |
| *Benthosema glaciale* | Glacier lanternfish | Myctophinae | Eastern Atlantic. Western Atlantic, Northwest Atlantic | fBenGla1.1 | https://doi.org/10.5281/zenodo.16941708 | CEES, University of Oslo | ~1.3 Gb | 22 | 26x |

**Table 2.**
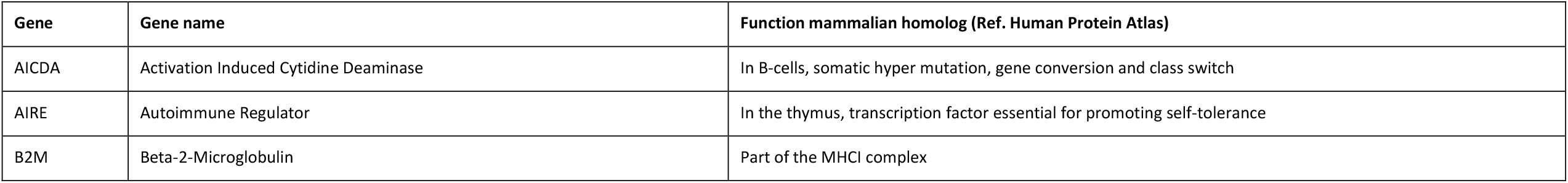
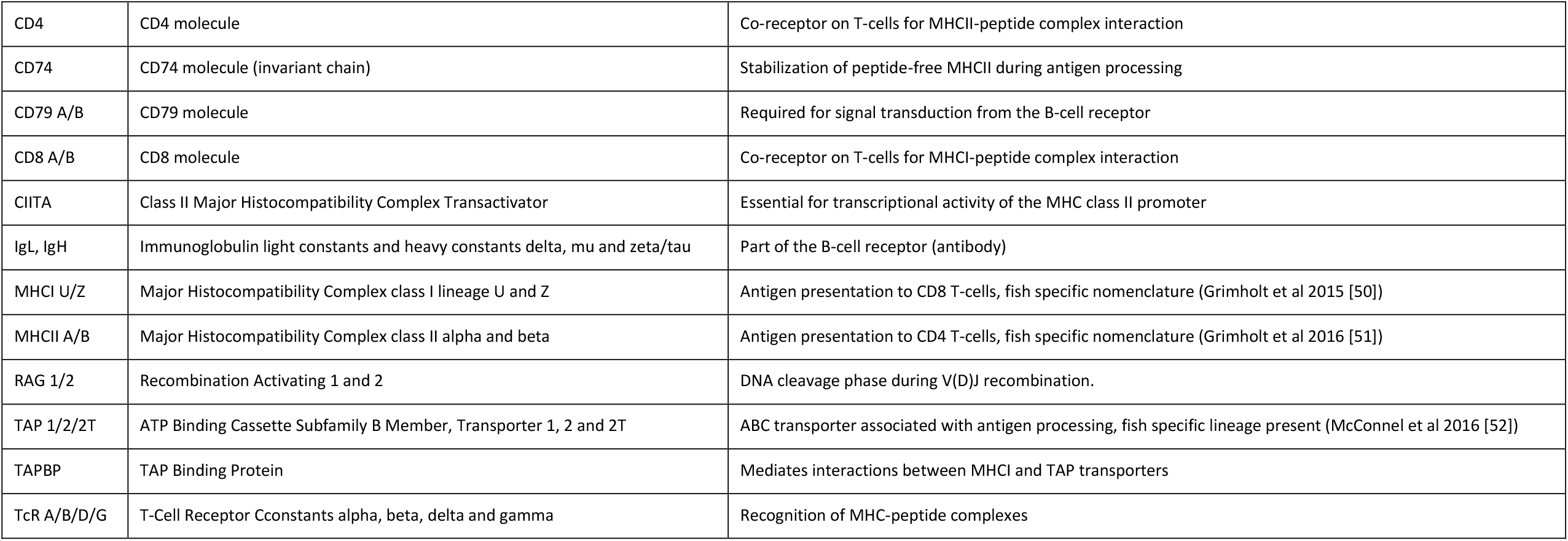
All immune-related genes manually annotated across genomes listed in Table 1.

With respect to antigen presentation all species showed intact representatives of at least one *MHCI* lineage and the MHCI co-receptor beta-2-microglobulin (*B2M*), but with large variation in gene copy numbers (**Figure 2 and 3, Table 3, Supplementary figure 2**). There were 2-6 copies of the non-classical Z lineage except for *B. pterotum* which has 27 copies. In contrast, the classic *MHCI U* lineage is absent from *Benthosema, Electrona* and *Protomyctophum*, but found in *Nannobrachium* and *Gymnoscopelus* with 16 to 45 gene copies (**Figure 3, Supplementary phylogenies**). Looking closer at the MHCI peptide loading machinery, we found that all species have intact tapasin (*TAPBP)*, transporter 1 ATP binding cassette subfamily B member (*TAP1)* and the teleost-specific *TAP2T*, but the loss of the classical *MHCI U* lineage coincides with a loss of the MHC-linked *TAP2* in *Benthosema, Protomyctophum* and *Electrona* (**Figure 2, Supplementary Figure 5**). *MHCIIA* and *MHCIIB* were found with less than 10 copies in *Benthosema, Electrona* and *Protomyctophum*. This is in stark contrast to *Nannobrachium* and *Gymnoscopelus* with closer to 200 copies (**Table 3**). However, only *Gymnoscopelus* and *Nannobrachium* have intact class II trans activator (*CIITA)* and invariant chain (*CD74)* which are required for *MHCII* expression and proper peptide loading in humans (**Figure 2**). Phylogenetic analysis shows two major clades (DA and DB, respectively) for both *MHCIIA* and *MHCIIB* sequences. Only *Gymnoscopelus* and *Nannobrachium* have gene copy representatives distributed across both clades in contrast to *Benthosema, Electrona* and *Protomyctophum* which are represented in DB only (**Supplementary phylogenies**).

**Figure 1.**
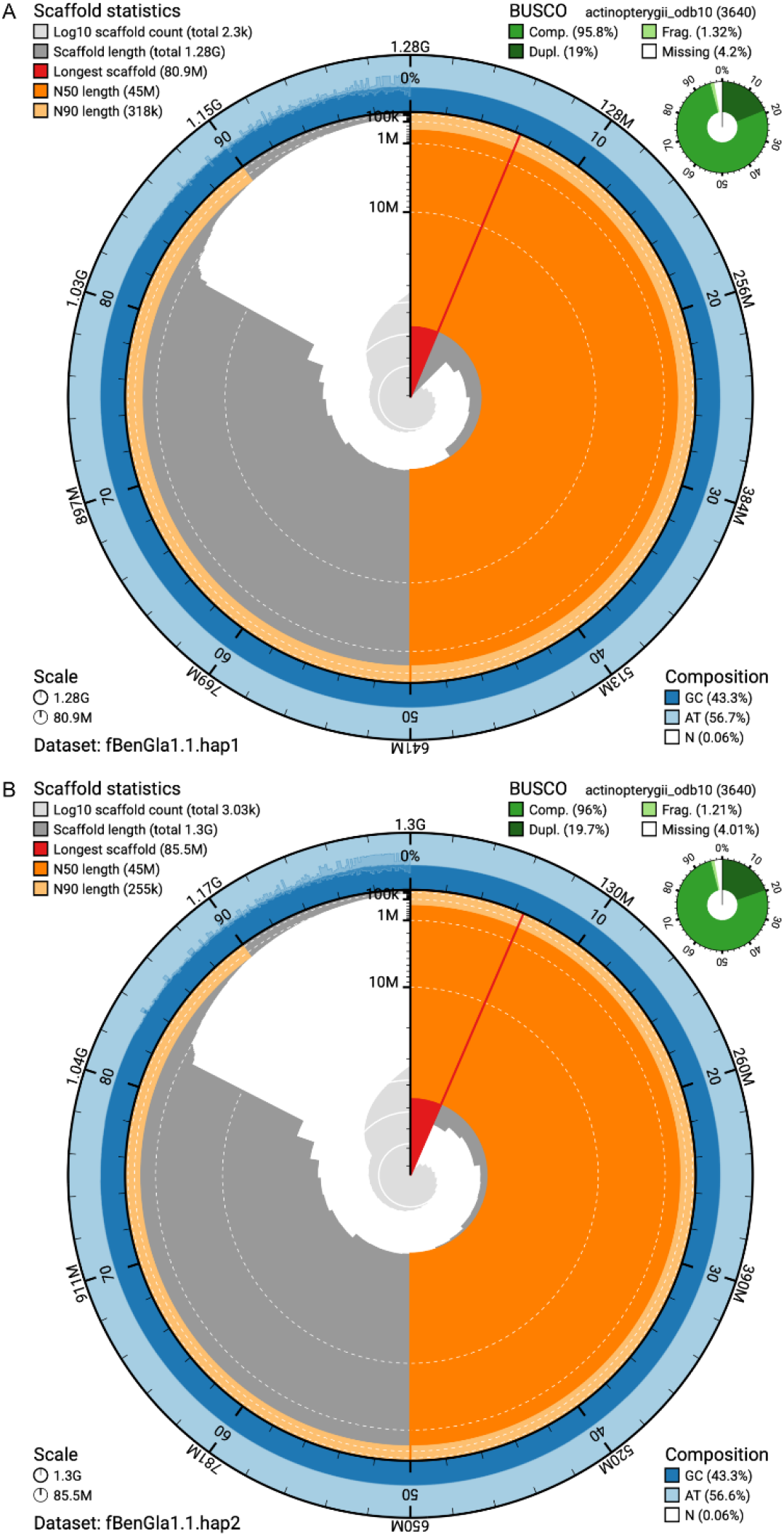
Metrics of the genome assemblies of Benthosema glaciale. The BlobToolKit Snailplots show N50 metrics and BUSCO gene completeness. The two outermost bands of the circle signify GC versus AT composition at 0.1% intervals. Light orange shows the N90 scaffold length, while the deeper orange is N50 scaffold length. The red line shows the size of the largest scaffold. All the scaffolds are arranged in a clockwise manner from the largest to the smallest and are shown in darker gray with white lines at different orders of magnitude. The light gray shows the cumulative scaffold count. The scale inset in the lower left corner shows the total amount of sequence in the whole circle, and the fraction of the circle encompassed in the largest scaffold.

**Figure 2.**
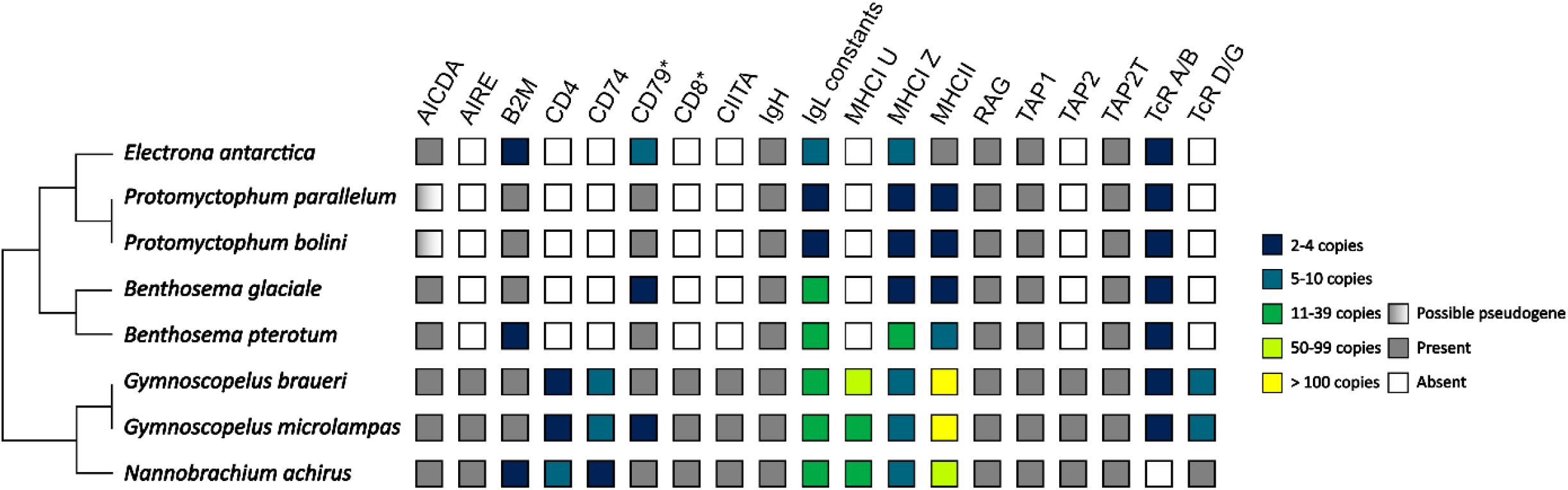
Overview of characterized adaptive immune genes projected onto a simplified Myctophidae phylogeny drawn after Denton et al. 2018 (note that several families have been omitted due to lack of genomic data representing these). White boxes are gene losses. Gradient white-grey boxes indicate possible pseudogenization. Grey boxes indicate gene presence with a single copy. Colored boxes indicate gene presence with gene expansion relative to the other species investigated (see color chart for copy numbers). *Highest copy number from alpha or beta subunit displayed. See Table 3 and Supplementary table 1 for details.

**Figure 3.**
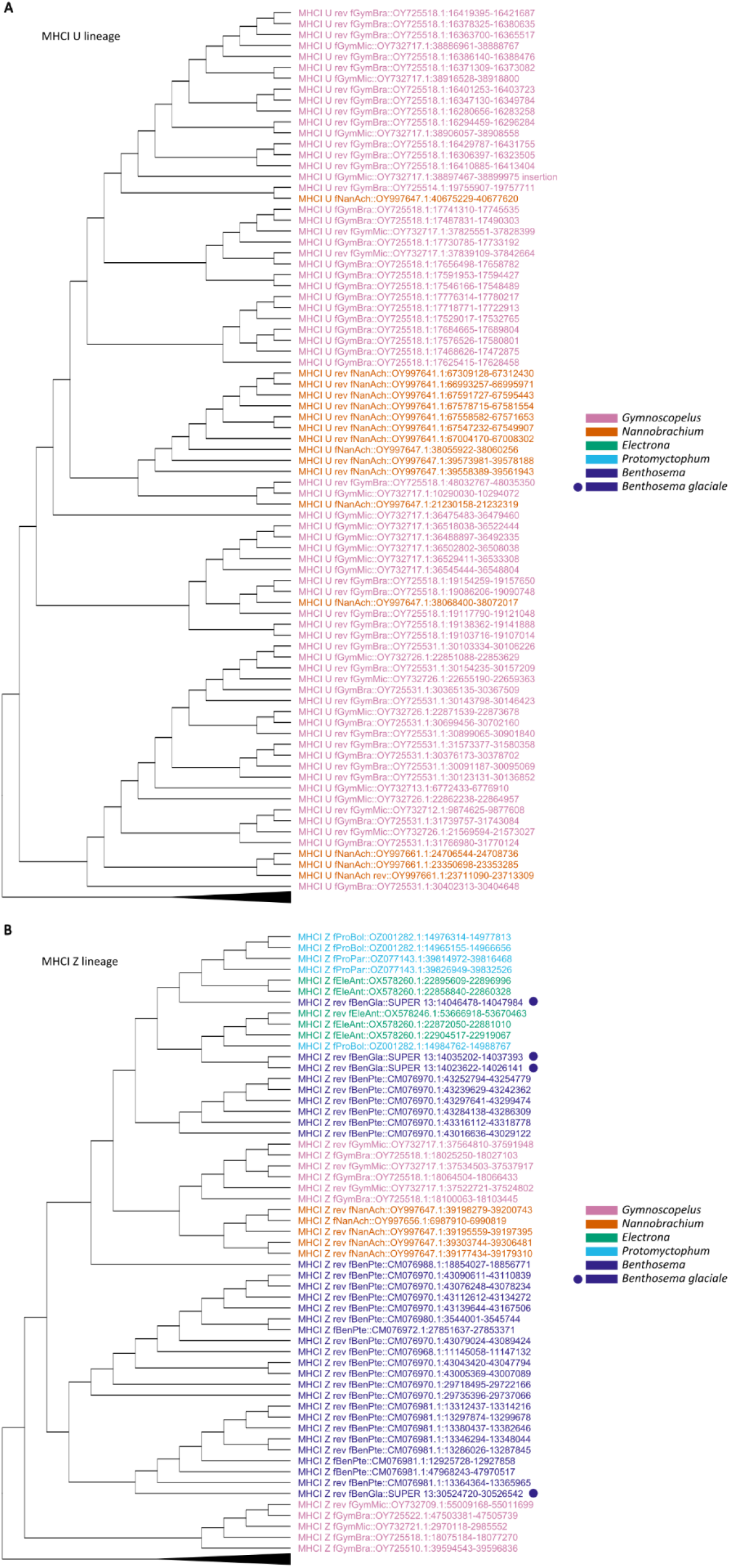
Subsets of the MHCI neighbor-joining tree on all MHCI protein sequences with Poisson substitution model, pairwise deletion and 500 bootstrap replicates made using MEGAX and shown as a cladogram. Reference sequences from Grimholt et al 2015 collapsed and colored in black. **A)** MHCI U lineage. **B)** MHCI Z lineage. Sequences colored by genus: Gymnoscopelus pink, Nannobrachium orange, Electrona green, Protomyctophum bright blue, Benthosema dark blue, Benthosema glaciale indicated by dot. Complete phylogeny in **Supplementary data**.

In terms of antigen recognition, all species have intact T-cell receptor loci containing 2-4 tandem copies of the *TcRB* constant and a single *TcRA* copy located elsewhere (except for *Nannobrachium* where no *TcRA* was found). *TcRD* and *TcRG* were only found in *Nannobrachium* and *Gymnoscopelus* and here located tandemly together with *TcRA* in *Gymnocopelus* (**Figure 2, Table 3, Supplementary figure 6**). The TcR co-receptors CD8 subunit alpha and beta (*CD8A*/*CD8B)* and CD4 subunit (*CD4)*, as well as the related transcription factor autoimmune regulator (*AIRE)* were found pseudogenized or lost from all genomes except in *Nannobrachium* and *Gymnoscopelus* (**Figure 2, Table 3, Supplementary figures 7-9**).

Finally, all species investigated have intact immunoglobulin heavy and light chain loci. For the heavy chain locus variation was observed with respect to the number of upstream variable regions as well as in the number of delta constants. The two *Gymnoscopelus* species also have a partial second locus consisting of only mu constants (**Figure 4**). The number of assembled light chain loci varied greatly between species ranging from 4 to 33 constant regions and 5 to 64 variable regions dispersed throughout the genome (**Table 3**). There appears to be reduced genetic diversity (low number of variable regions) in *Protomyctophum* and *Electrona*, intermediate diversity for *Benthosema* and high diversity for *Nannobrachium* and *Gymnoscopelus* (**Table 3**). All species have present CD79 alpha and beta (*CD79 a/b)* required for signaling through the membrane bound immunoglobulin B-cell receptor. Furthermore, all species have intact recombination activating 1 and 2 (*RAG1/2)* required for the recombination of both immunoglobulins and T-cell receptors as well as activation induced cytidine deaminase (*AICDA)* required for class-switch recombination and somatic hypermutation in humans. However, *AICDA* in the two *Protomyctophum* species was of notable poorer sequence integrity indicating possible pseudogenization or assembly issues (**Figure 2, Supplementary figures 10–12, Supplementary data 1**).

**Table 3.** Fully assembled adaptive immune genes located to scaffolds in the investigated genomes. Details listed in Supplementary data 1.

| Species | IgH variable | IgH constants | IgL variable | IgL constants | TcR variable | TcR constants | MHCI U | MHCI Z | MHCII A | MHCII B |
| --- | --- | --- | --- | --- | --- | --- | --- | --- | --- | --- |
| *Benthoosema glaciale* | 17 | IgM, IgD | 32 | 24 | 15 | TcRA, TcRB | 0 | 4 | 2 | 2 |
| *Benthoosema pterotum* | 32 | IgM, IgD | 38 | 32 | 27 | TcRA, TcRB | 0 | 27 | 5 | 5 |
| *Electrona antarctica* | 29 | IgM, IgD | 17 | 9 | 10 | TcRA, TcRB | 0 | 5 | 1 | 1 |
| *Protomyctophum bolini* | 7 | IgM, IgD | 6 | 4 | 4 | TcRA, TcRB | 0 | 3 | 2 | 2 |
| *Protomyctophum parallelum* | 6 | IgM, IgD | 5 | 4 | 4 | TcRA, TcRB | 0 | 2 | 4 | 4 |
| *Nannobranchium achirus* | 42 | IgM, IgD | 64 | 33 | 18 | TcRB, TcRD,<br>TcRG | 16 | 5 | 83 | 92 |
| *Gymnoscopelus microlampas* | 48 | IgM, IgD | 61 | 32 | 89 | TcRA, TcRB,<br>TcRD, TcRG | 20 | 5 | 25 | 192 |
| *Gymnoscopelus braueri* | 66 | IgM, IgD | 57 | 32 | 89 | TcRA, TcRB,<br>TcRD, TcRG | 45 | 6 | 21 | 195 |

**Figure 4.**
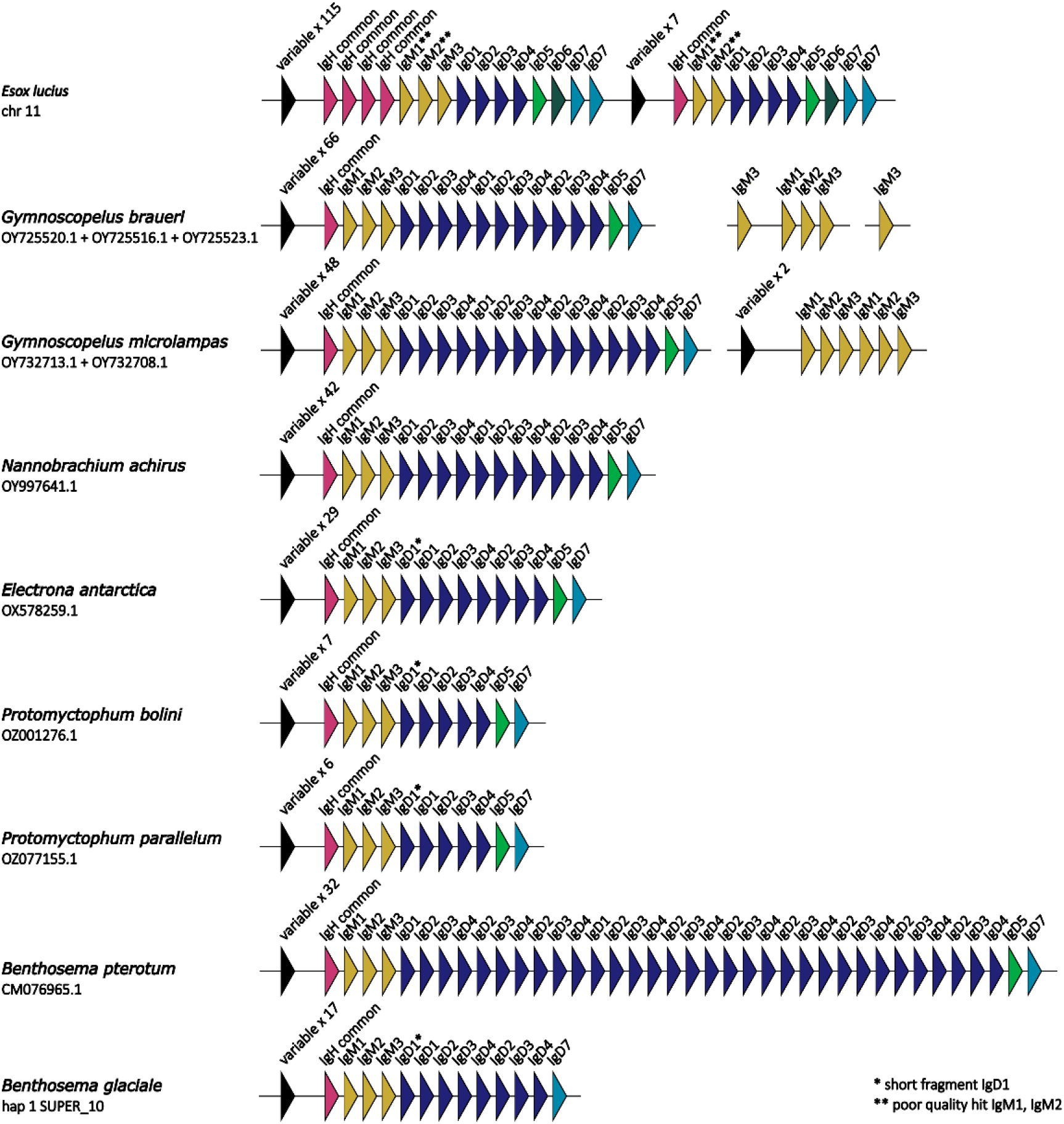
Manually annotated immunoglobulin heavy chain loci in all investigated species. Annotation of constant domains based on the IgH locus of Northern pike (Esox lucius) [94, 95]. The IgH common constant (present in both in IgM and IgD transcripts) is highlighted in crimson. IgM constants are mustard. IgD constants 1-4 are dark blue. IgD constants 5-7 are shown in light green, dark green and light blue, respectively. Variable region in black with copy number written above. Joining regions not shown. IgT/Z not drawn due to no BLAST hits in the Myctophidae genomes.

## Discussion

Here we present a chromosome level genome assembly of the Glacier lanternfish *B. glaciale*. Compared to other available Myctophidae genomes it is similar in size but with a slightly reduced number of chromosomes (22 vs. 24) (**Table 1**). The difference in autosome number could be related to the significant diversity observed within the Myctophidae demonstrated by e.g. uncertain placement of the two *Benthosema* species in phylogenetic studies [25, 28]. It is also well documented that chromosome number may vary extensively in closely related teleost species such as in codfishes (Gadidae) where Atlantic cod has 23 chromosomes compared to 18 in polar cod (*Boreogadus saida*) and 15 in Arctic cod (*Arctogadus glacialis*) [41]. The *B. glaciale* genome is also comparable to the *Electrona antarctica (E*.*antarctica)* and *B. pterotum* genomes with respect to completeness with more than 95 % complete BUSCOs. They also show between 9 and 15 % duplicated BUSCOs compared to 19 % duplicated shown here (**Figure 1, Table 1**) [42, 43]. This level of duplication is higher than many other teleost genomes (e.g. 1.8 % in European smelt (*Osmerus eperlanus*) [44], 2.4% in the greater Argentine (*Argentina silus*) [45], and 0.9% in Atlantic cod (*Gadus morhua*) [46]. Thus, the duplicated BUSCOs in *Benthosema* and *Electrona* assemblies may be the result of an lineage-specific event rather than the ancient teleost whole genome duplication (3R WGD) dated to around 320 Mya [47]. Overall, the generated *B. glaciale* genome is of comparative quality and completeness to other related genomes.

Studies in some other species-rich teleost orders have reported innovative immune strategies that deviate from what has long been perceived as the vertebrate consensus with respect to adaptive immunity [48]. This includes loss of *MHCII* in the entire codfish order (Gadiformes) [49–54], in some pipefish (Syngnathiformes) clades [55] and in some anglerfish (Lophiiformes) clades [56, 57]. These findings represent challenges for our biological and immunological understanding and may consequently affect knowledge-based management efforts.

Our results show that the clade represented by *Benthosema, Electrona* and *Protomyctophum* has lost what is regarded as the classical MHC class I lineage in teleosts (MHCI U, [58]) (**Figure1 and 2, Table 3**) as well as the required co-receptor CD8 [59] (**Figure1, Table 3**). These findings corroborate earlier implications regarding MHCI function in *B. glaciale* reported by Malmstrøm *et al*. where the low coverage sequencing strategy did not allow for further in-depth analysis [54]. With respect to MHCII, we observe a reduced *MHCII* repertoire in *Benthosema, Electrona* and *Protomyctophum* (**Table 3**).

Phylogenetic analyses demonstrate that Myctophidae *MHCII* genes segregate into two major clades similar to that shown in Dijkstra *et al*. [62]. *Gymnoscopelus* and *Nannobrachium* genes are distributed throughout both the classical DA and non-classical DB groups. In contrast, *Benthosema, Electrona* and *Protomyctophum MHCII* genes are only found non-classical DB (**Supplementary phylogenies**).

The simultaneous absence of central co-factors required for classical MHCII functionality such as *CD74* [60], *CIITA* [61] and *CD4* [59] (**Figure 2**) indicates loss of classical MHCII functionality [62, 63]. Loss of MHCII has previously been reported in codfishes [49, 54], in some pipefishes [55] and in some anglerfishes [57]. However, those species do not have any remnants of *MHCII* in their genomes [49, 54, 55] in contrast to our findings in *Benthosema, Electrona* and *Protomyctophum* (**Table 3**).

All in all, we find that *Benthosema, Electrona* and *Protomyctophum* have lost all classical MHC functionality from both the class I and class II systems. Loss of both MHC systems has previously been implied in some anglerfish species through the lack of required co-factors [56]. However, there are only incomplete genomes available for these species leaving the question regarding their MHC status. To our knowledge, our study is the first to firmly demonstrate an alternative adaptive immune strategy with loss of all classic MHC (I and II) function while retaining *MHC* gene copies in the genome. Further investigations are needed to evolutionary time the event(s) and in-depth elucidate the function of the non-classical MHCs present in the genomes of *Benthosema, Electrona* and *Protomyctophum*.

With the loss of classic MHC function, one might also expect a loss of corresponding T-cell function. *Benthosema, Electrona* and *Protomyctophum* all have intact alpha and beta T-cell receptor loci - albeit with fewer variable regions compared to *Gymnoscopelus* (**Figure 2, Table 3**). In terms of function, the alpha-beta T-cells are the most abundant and best described T-cell populations, where the majority interact with peptides presented on MHCs. These conventional receptors are restricted by MHC because the ligands presented have to fit within the MHC pocket [64]. Conventional peptide-MHC recognizing T-cell receptors also undergo thymic selection in mammals to ensure that they react within an acceptable framework – a process that requires the AIRE transcription factor [65].

Fish do have a thymic structure [66], but the loss of AIRE in *Benthosema, Electrona* and *Protomyctophum* indicates that their T-cell receptors do not go through thymic selection (**Figure 2, Table 3**). There are reports on unconventional alpha-beta T-cell receptors in some species (e.g. humans, mice and xenopus) recognizing non-polymorphic antigens [65, 67, 68] - possibly an explanation for the present alpha-beta T-cell receptor loci in *Benthosema, Electrona* and *Protomyctophum*. Loss of alpha-beta T-cell receptors has been observed in some anglerfish [56]. To our knowledge, our study is the first to demonstrate loss of classic T-cell receptor function while retaining intact alpha-beta T-cell receptor loci in the genome, and further investigations are needed to elucidate the function of T-cell receptors in an MHC-impaired environment.

*Benthosema, Electrona* and *Protomyctophum* have also lost the gamma and delta T-cell receptor loci (**Figure 2**). Delta and alpha constants are tightly linked in humans with overlapping variable regions [69] which is also the case for *Gymnoscopelus* (**Supplementary** data 1). Gamma-delta T-cells are typically unrestricted by MHC [64], and in humans they are suspected to recombine during development, colonize tissues and then regulate homeostasis and immune surveillance where they reside [65]. As gamma-delta T-cell receptors are unrestricted by MHC it remains elusive why they have been lost from *Benthosema, Electrona* and *Protomyctophum*. Our findings warrant additional functional studies to understand gamma-delta T-cells in lanternfishes and teleosts in general.

All investigated Mytan;ledae species, including Myctophinae (*Benthosema, Electrona, Protomyctophum*) with loss of classical MHC functionality, should be able to make natural antibodies based on the presence of both immunoglobulin light and heavy chain loci and cofactors needed for V(D)J recombination and downstream signaling [70] (**Figure 1** and **2**). The possibility of generating diversity beyond the germline is also indicated by the presence of *AICDA* needed for somatic hyper mutation [70] (**Figure 2**). A similar situation is found in Atlantic cod and codfishes where the lack of *MHCII* could affect the production of antibodies. Also here an intact immunoglobulin locus as well as *AICDA* is found [53, 71]. However, in-depth studies of codfish AICDA demonstrate that the enzyme is inactive with respect to somatic hypermutation [53]. Studies of immunoglobulin diversity in unchallenged Atlantic cod demonstrates comparable diversity to that of other teleosts - tentatively connected to the triplication of the heavy chain locus in this species [71]. Lanternfish will require further studies to fully understand the immunoglobulin diversity in these species and to what degree diversity beyond the germline is occurring.

In sum, we find that there are (at least) two vastly different adaptive immune strategies within Myctophiformes. The Myctophinae (*Benthosema, Electrona* and *Protomyctophum*) have lost all classical MHC functionality for both the class I and class II systems but retain non-classical MHC and T-cell receptor genes as well as intact immunoglobulin loci. In contrast, *Gymnoscopelus* and *Nannobrachium* appear to have maintained a conventional adaptive immune system with slight differences in the degrees of available genetic diversity (**Figure 2, Figure 3, Figure 4, Table 3**). Our findings thus raise new questions regarding the adaptive immune system in lanternfishes and in teleosts in general. Further studies are needed to elucidate the function of non-classical adaptive immune genes such as MHCI Z and MHCII DB. Interesting contrasts can also be made between Myctophidae where function is lost through the removal of crucial co-factors compared to losses of key genes in e.g. codfishes and pipefishes. Our study also demonstrates how crucial obtaining complete genomes are to obtain optimal biological understanding.

## Methods

### Sample acquisition, DNA isolation, sequencing and assembly

One individual of *Benthosema glaciale* was obtained by shrimp trawl in Trondheimsfjorden 1^st^ of July 2021 – position 63.717229 - 9.875115 at 219 m depth (**Supplementary figure 13**).

The individual was put on ethanol and later shipped to the University of Oslo. Here, the individual was evaluated a second time with respect to species identification. The sample was partially dissected and DNA isolated from tail/muscle, liver/internal organs and heart/brain/skin.

All DNA isolation, library preparation and sequencing was performed by the Norwegian Sequencing Centre. In short, DNA isolation was performed using Circulomics Nanobind BIG DNA prep tissue protocol according to the manufacturer’s instructions. The liver/internal organs sample was selected for library preparation due to high yield of DNA and the best overall fragment length distribution (**Supplementary table 1 and Supplementary figure 14-15**).

Library preparation was performed using the Pacific Biosciences protocol for HiFi library prep SMRTbell® ExpressTemplate Prep Kit 2.0. The DNA was fragmented to 15-20 kb fragments using Megaruptor 3 (Diagenode) and the final library was size selected using BluePippin (Sage Science Inc) with a 8kb cut-off. The library was sequenced on three 8M SMRT cells on Sequel II (PacBio) using the Sequel II Binding Kit 2.2 and Sequencing chemistry v2.0, adaptive loading and movie time 30 hours. The last two SMRT cells were sequenced using 2 hours pre-extension. Circular consensus sequences (CCS) were generated using the CCS pipeline (SMRT Link v10.1.0.119588 SMRTcell 1 and v10.2.0.133434 SMRTcell 2 and 3) with default settings (**Supplementary table 2**).

For the HiC library preparation we used 115 mg of ethanol-preserved muscle tissue with the Arima High Coverage HiC kit (November 2021) according to the manufacturer’s instructions for animal tissues with standard input tissue amounts (Arima Genomics Inc). The final library was sequenced on the Illumina Novaseq 6000 with 2×150 bp paired end mode (**Supplementary table 2**).

### Assembly

A full list of relevant software tools and versions is presented in **Supplementary table 3**. KMC was used to count k-mers of size 32 in the PacBio HiFi reads, excluding k-mers occurring more than 10,000 times [72]. GenomeScope was run on the k-mer histogram output from KMC to estimate genome size, heterozygosity and repetitiveness while ploidy level was investigated using Smudgeplot [73]. HiFiAdapterFilt was applied on the HiFi reads to remove possible remnant PacBio adapter sequences [74]. The filtered HiFi reads were assembled using hifiasm with Hi-C integration resulting in a pair of haplotype-resolved assemblies, pseudo-haplotype one (hap1) and pseudo-haplotype two (hap2) [75]. Unique k-mers in each assembly/pseudo-haplotype were identified using meryl and used to create two sets of Hi-C reads, one without any k-mers occurring uniquely in hap1 and the other without k-mers occurring uniquely in hap2 [76]. K-mer filtered Hi-C reads were aligned to each scaffolded assembly using BWA-MEM with -5SPM options [77]. The alignments were sorted based on name using samtools before applying samtools fixmate to remove unmapped reads and secondary alignments and to add mate score, and samtools markdup to remove duplicates [78]. The resulting BAM files were used to scaffold the two assemblies using YaHS with default options [79]. FCS-GX was used to search for putative contamination [80]. Contaminated sequences were removed. The mitochondrion was searched for in reads using Oatk [81]. Merqury was used to assess the completeness and quality of the genome assemblies by comparing to the k-mer content of the Hi-C reads [76]. BUSCO was used to assess the completeness of the genome assemblies by comparing against the expected gene content in the actinopterygii lineage [82]. Gfastats was used to output different assembly statistics of the assemblies [83]. The assemblies were manually curated using PretextView and Rapid curation 2.0. Chromosomes were identified by inspecting the Hi-C contact map in PretextView. BlobToolKit and BlobTools2 in addition to blobtk were used to visualize assembly statistics and GC-coverage plots [84, 85]. To generate the Hi-C contact map image, the Hi-C reads were mapped to the assemblies using BWA-MEM [77] using the same approach as above, before PretextMap was used to create a contact map which was visualized using PretextSnapshot.

We annotated the genome assemblies using a pre-release version of the EBP-Nor genome annotation pipeline (https://github.com/ebp-nor/GenomeAnnotation). First, AGAT (https://zenodo.org/record/7255559) agat_sp_keep_longest_isoform.pl and agat_sp_extract_sequences.pl were used on the GRCz11 genome assembly and annotation to generate one protein (the longest isoform) per gene. Miniprot was used to align the proteins to the curated assemblies [86]. UniProtKB/Swiss-Prot (UniProt Consortium 2023) release 2023_03 in addition to the vertebrata part of OrthoDB v11 were also aligned separately to the assemblies [87]. Red was run via redmask (https://github.com/nextgenusfs/redmask) on the assemblies to mask repetitive areas [88]. GALBA was run with the GRCh38 proteins using the miniprot mode on the masked assemblies [86, 89–92]. The funannotate-runEVM.py script from Funannotate was used to run EvidenceModeler on the alignments of GRCz11 proteins, UniProtKB/Swiss-Prot proteins, vertebrata proteins and the predicted genes from GALBA [20]. The resulting predicted proteins were compared to the protein repeats that Funannotate distributes using DIAMOND blastp and the predicted genes were filtered based on this comparison using AGAT. The filtered proteins were compared to the UniProtKB/Swiss-Prot release 2023_03 using DIAMOND blastp to find gene names and InterProScan was used to discover functional domains [90]. AGATs agat_sp_manage_functional_annotation.pl was used to attach the gene names and functional annotations to the predicted genes. EMBLmyGFF3 was used to combine the fasta files and GFF3 files into a EMBL format for submission to ENA [93].

### Gene identification

For comparative purposes additional lanternfish genomes were downloaded from NCBI in addition to the *Benthosema glaciale* genome generated here (**Table 1**).

Query protein sequences for central immune genes were obtained from NCBI or Ensembl covering the genes listed in **Table 2**. Fully assembled genes placed to a chromosome were included. For suspected gene losses all unplaced scaffolds were also included in the search, and local gene synteny used to verify the gene loss or pseudogene when possible.

## Supporting information

Supplementary_phylogenies

## Acknowledgement

The authors would like to thank Professor Stein Kaartvedt University of Oslo for help with species identification. This project received data management and infrastructure support from ELIXIR Norway, supported by the Research Council of Norway’s grant 270068, the University of Bergen, the University of Oslo, the Arctic University of Norway in Tromsø, the Norwegian University of Science and Technology and the Norwegian University of Life Sciences: NMBU. The authors acknowledge support from the National Infrastructure for High Performance Computing and resources provided by Sigma2 as well as Data Storage in Norway (project NN9244 and NN8013K) for computational work.

We would also like to thank the Norwegian Sequencing Centre (NSC: https://www.sequencing.uio.no) for their sample preparation and sequencing services, and the crew of RV Gunnerus (NTNU). The project was funded by the S. G. Sønneland Foundation to MHS and the University of Oslo Life Sciences Convergence programme to the project “Comparative immunology of fish and humans (COMPARE)” and by the Research Council of Norway project 326819 (The Earth Biogenome Project Norway; EBP-Nor) to KSJ.

## Data availability

The genome assemblies and gene annotations are available at https://doi.org/10.5281/zenodo.16941708.

