## Supplementary_information for "The genome of the Glacier lanternfish shows loss of MHC I and II function and provides insight into evolution of lanternfish immune systems"

### Supplementary figures and tables

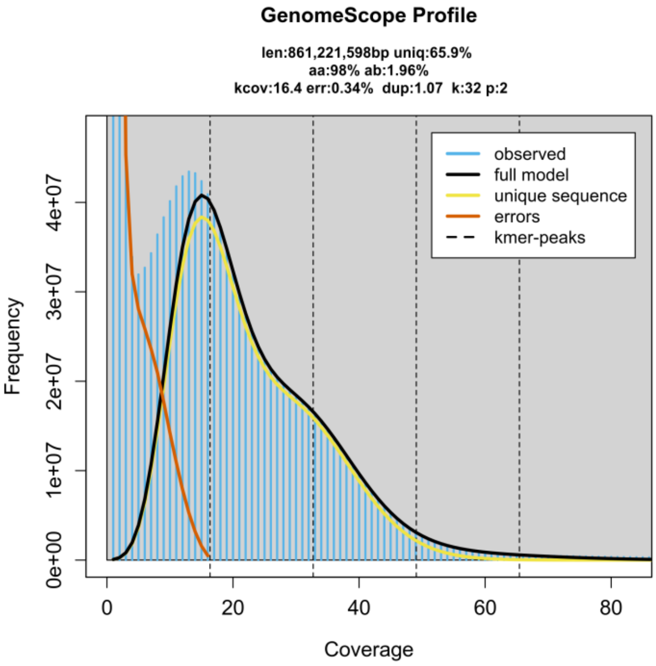


*Supplementary figure 1 GenomeScope profile of the HiFi reads from the sequenced individual. This analysis estimates a 861 Mb genome, with 1.96 % heterozygosity.*

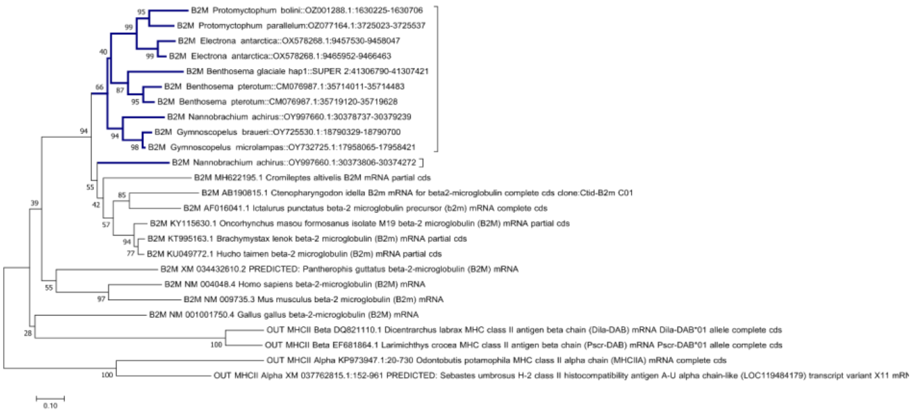


*Supplementary figure 2 Neighbor-joining tree on B2M protein sequences with Poisson substitution model, pairwise deletion and 500 bootstrap replicates made using MEGAX. Myctophiformes species in blue.*

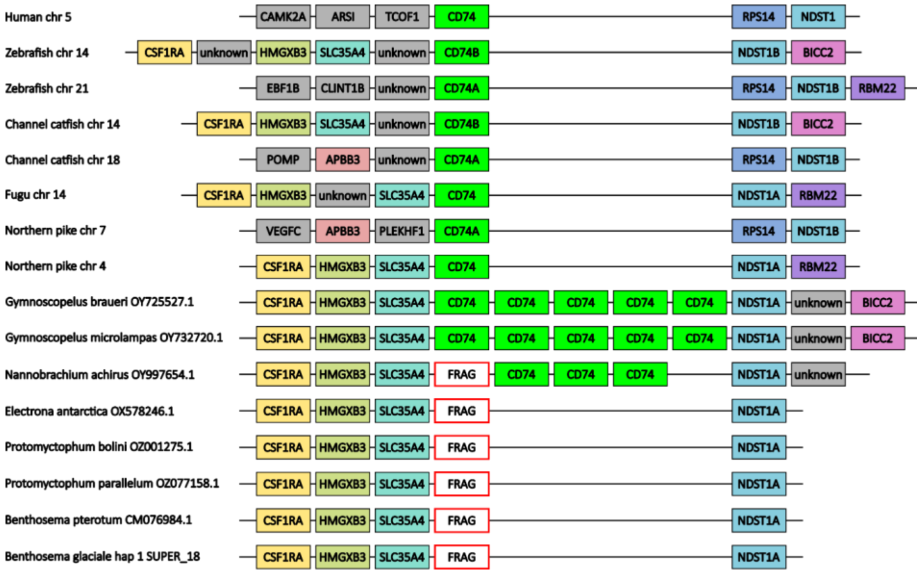


*Supplementary figure 3 Local gene synteny drawn based on BLAST results in the Myctophiformes genomes and gene annotations from Ensembl for the remaining species. CD74 expansions detected in Gymnoscopelus braueri, Gymnoscopelus microlampas and Nannobrachium achirus. Fragments detected for CD74 in Electrona antarctica, Protomyctophum bolini, Protomyctophum parallelum, Benthosema pterotum and Benthosema glaciale.*

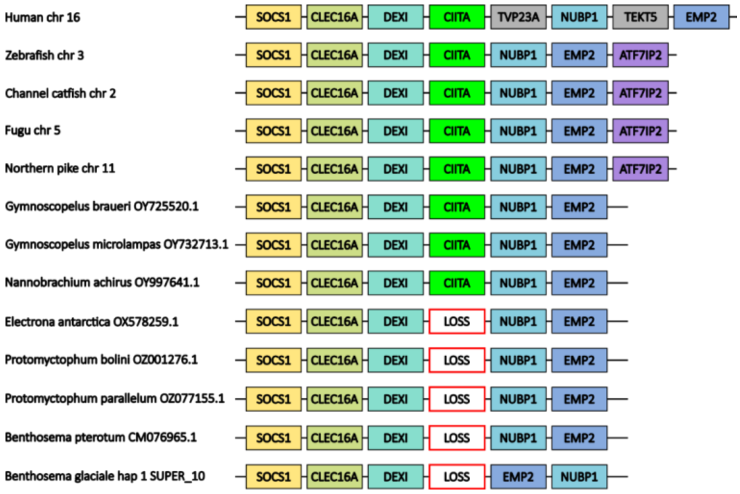


*Supplementary figure 4 Local gene synteny drawn based on BLAST results in the Myctophiformes genomes and gene annotations from Ensembl for the remaining species. Gene loss detected for CIITA in Electrona antarctica, Protomyctophum bolini, Protomyctophum parallelum, Benthosema pterotum and Benthosema glaciale.*

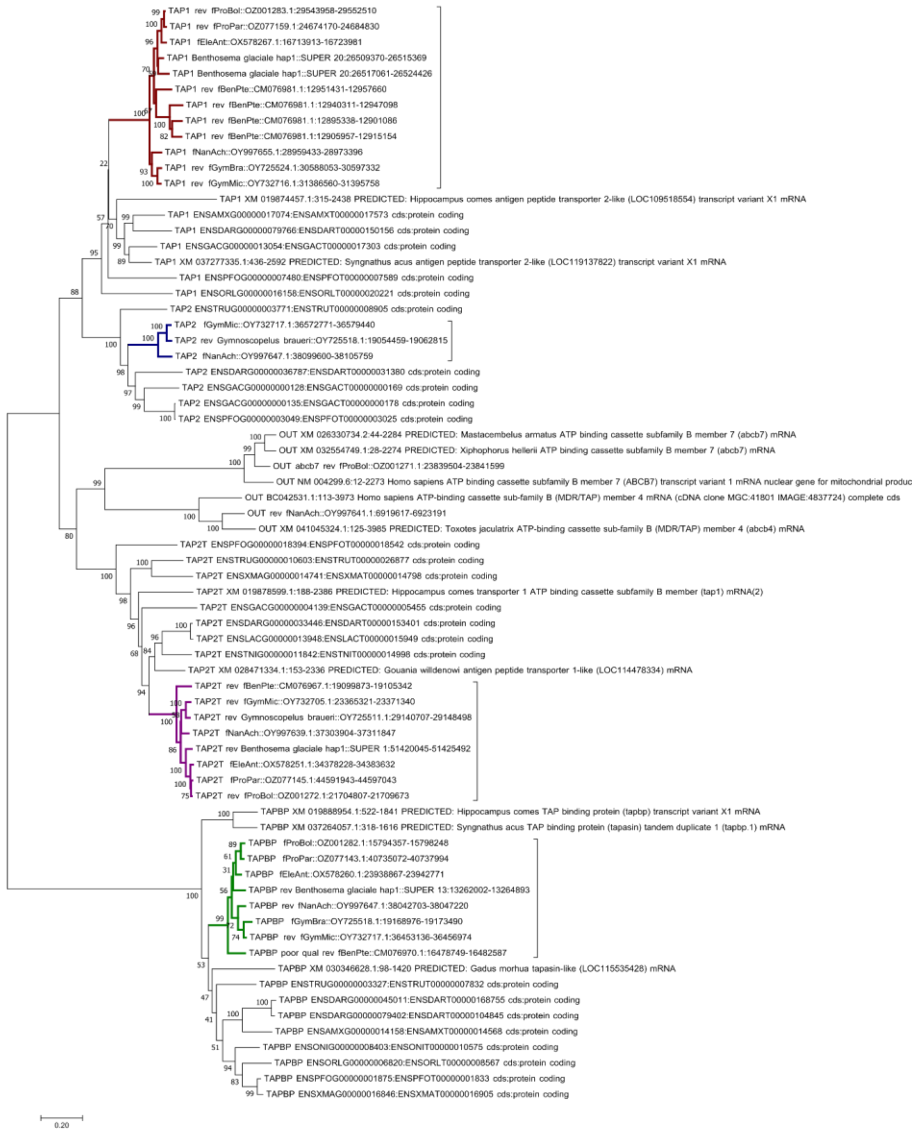


*Supplementary figure 5 Neighbor-joining tree on TAPBP, TAP1, TAP2 and TAP2T protein sequences with Poisson substitution model, pairwise deletion and 500 bootstrap replicates made using MEGAX. Myctophiformes species in red (TAP1), blue (TAP2), purple (TAP2T), and green (TAPBP).*

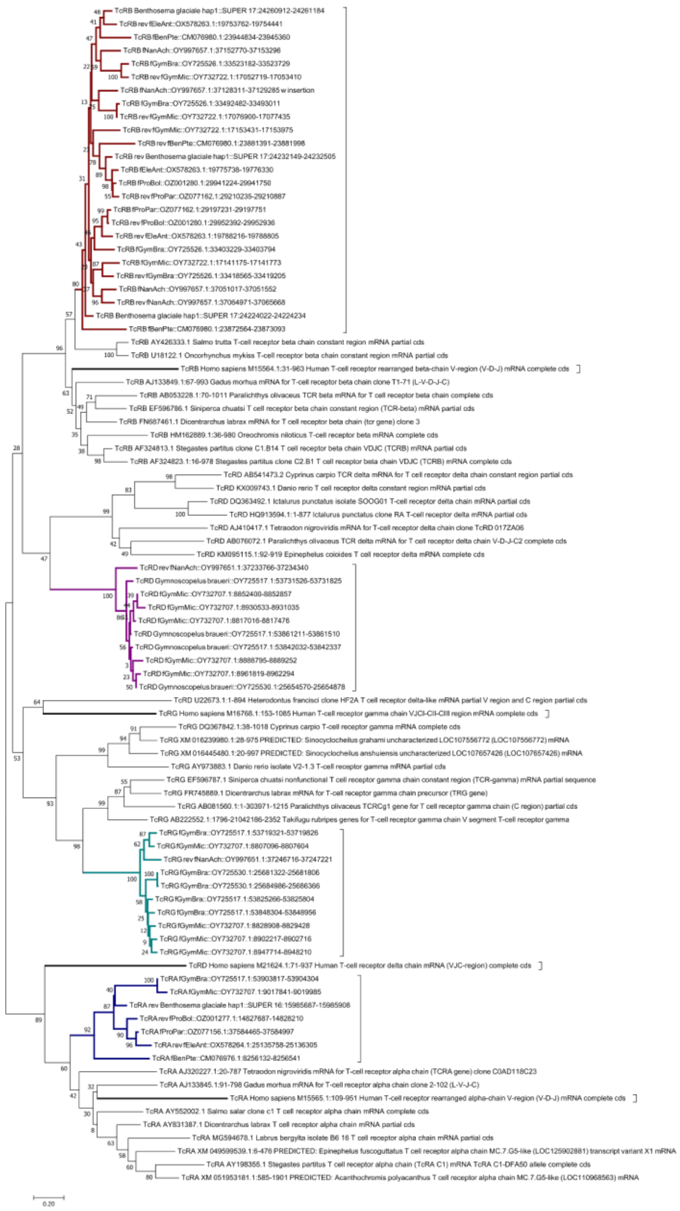

*Supplementary figure 6 Neighbor-joining tree on TcR constant region protein sequences with Poisson substitution model, pairwise deletion and 500 bootstrap replicates made using MEGAX. Myctophiformes species in blue (TcRA), red (TcRB), purple (TcRD) and teal (TcRG). Human representatives in bold black.*

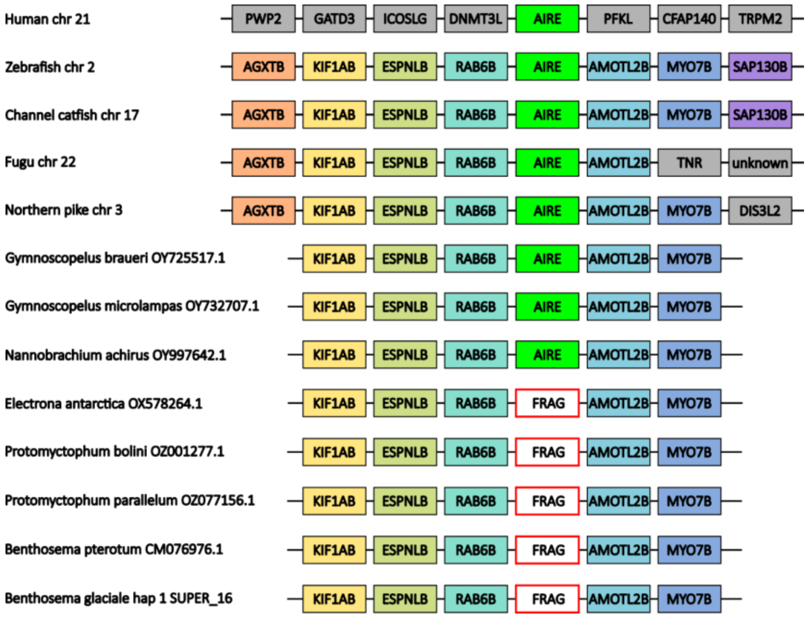


*Supplementary figure 7 Local gene synteny drawn based on BLAST results in the Myctophiformes genomes and gene annotations from Ensembl for the remaining species. Fragments detected for AIRE in Electrona antarctica, Protomyctophum bolini, Protomyctophum parallelum, Benthosema pterotum and Benthosema glaciale.*

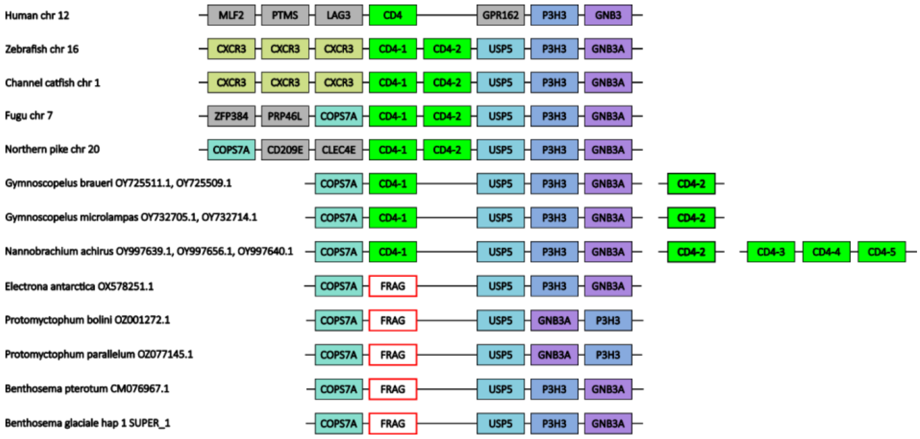


*Supplementary figure 8 Local gene synteny drawn based on BLAST results in the Myctophiformes genomes and gene annotations from Ensembl for the remaining species. Fragments detected for CD4 in Electrona antarctica, Protomyctophum bolini, Protomyctophum parallelum, Benthosema pterotum and Benthosema glaciale.*

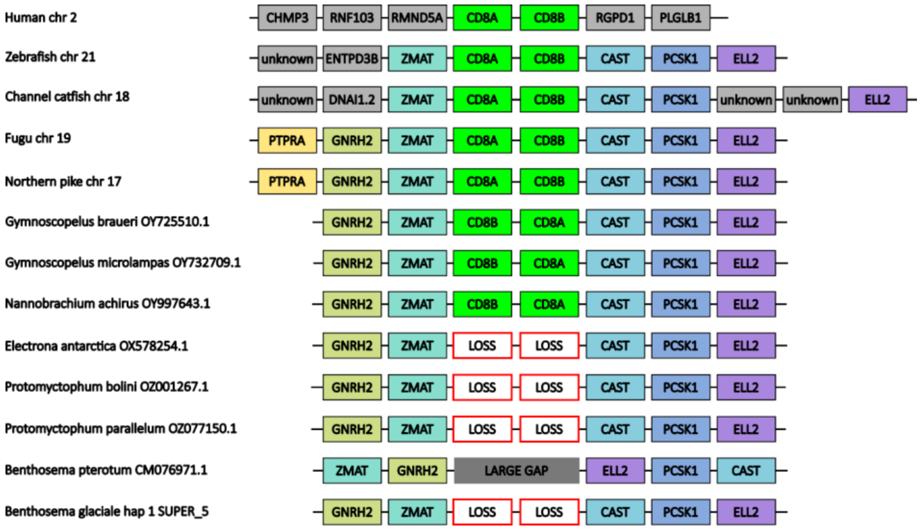


*Supplementary figure 9 Local gene synteny drawn based on BLAST results in the Myctophiformes genomes and gene annotations from Ensembl for the remaining species. Loss of both CD8 subunits in Electrona antarctica, Protomyctophum bolini, Protomyctophum parallelum and Benthosema glaciale. Assembly gap in syntenic region for Benthosema pterotum.*

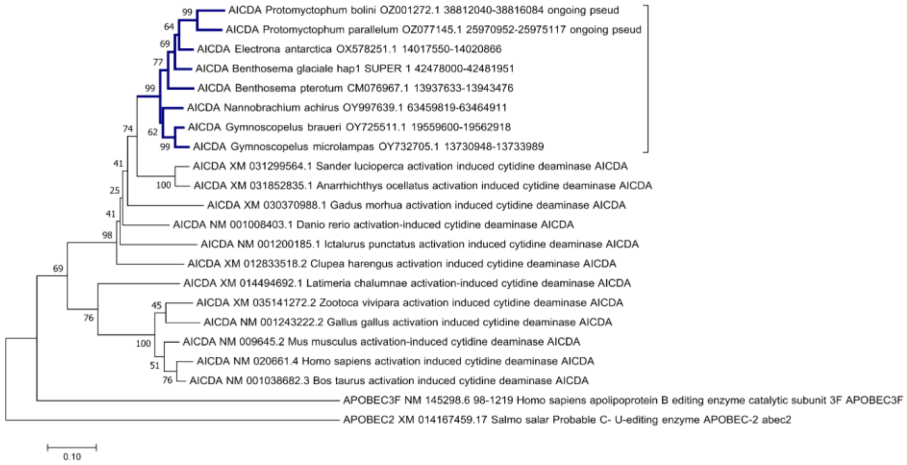


*Supplementary figure 10 Neighbor-joining tree on AICDA protein sequences with Poisson substitution model, pairwise deletion and 500 bootstrap replicates made using MEGAX. Myctophiformes species in blue.*

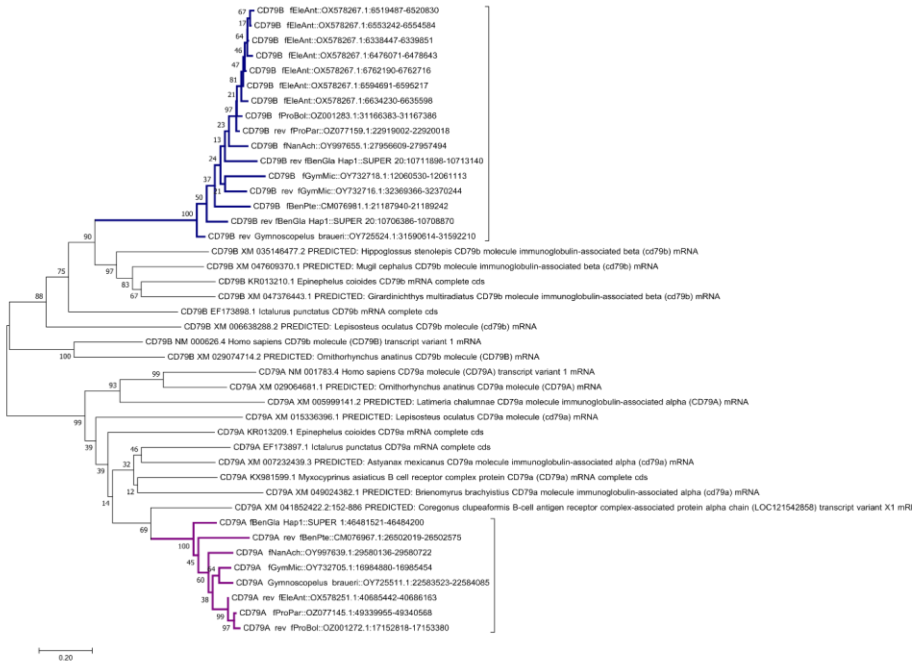


*Supplementary figure 11 Neighbor-joining tree on CD79 alpha and beta subunit protein sequences with Poisson substitution model, pairwise deletion and 500 bootstrap replicates made using MEGAX. Myctophiformes species in blue and purple.*

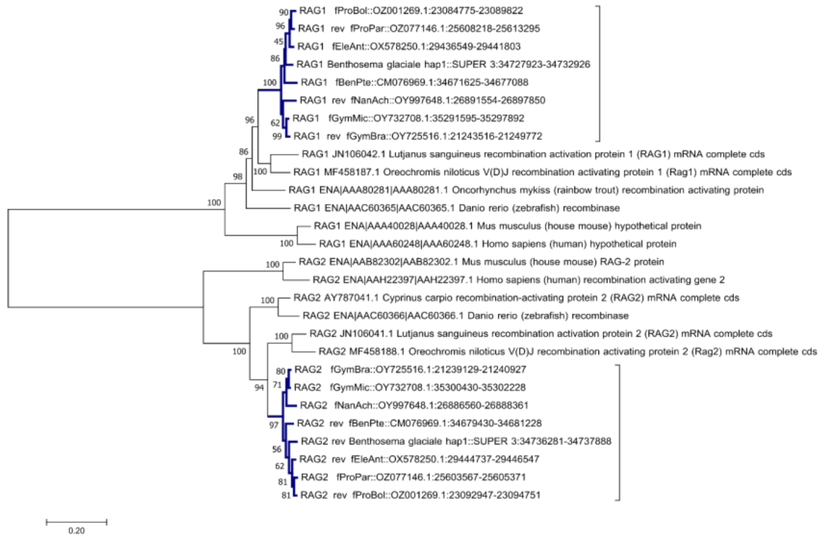


*Supplementary figure 12 Neighbor-joining tree on RAG1 and RAG2 protein sequences with Poisson substitution model, pairwise deletion and 500 bootstrap replicates made using MEGAX. Myctophiformes species in blue.*

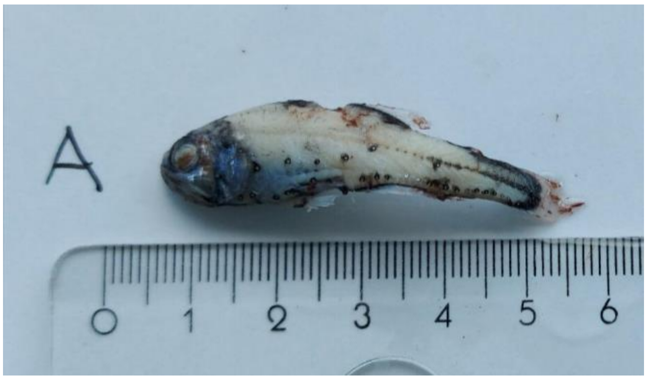


*Supplementary figure 13 Picture of the collected sample post ethanol conservation displayed with a centimeter ruler.*

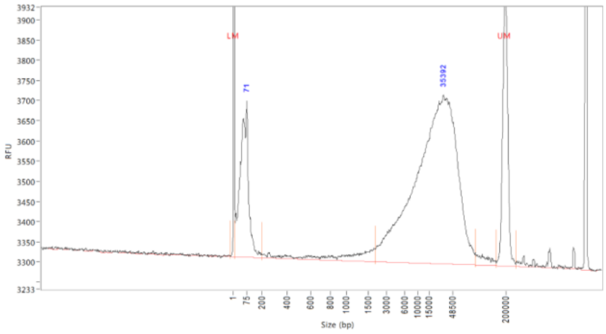


*Supplementary figure 14* Fragment analyzer trace of sample BenGla-3 used in library preparation.

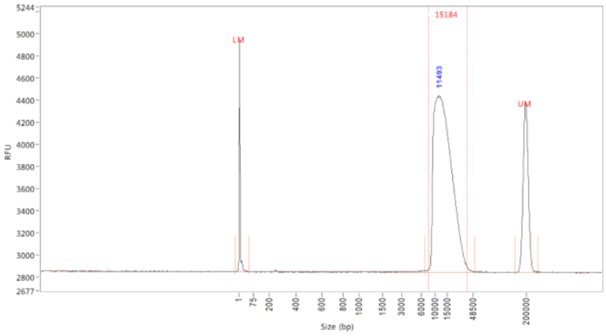


*Supplementary figure 15* Final size selected library fragment trace of sample BenGla-3 after SMRTbell ExpressTemplate prep kit, fragmentation (Megaruptor) and size selection (Bluepippin).

*Supplementary table 1* Results from Circulomics Nanobind DNA isolation protocol. Concentration was determined by the average of 3 measurements using Qubit BR

| Sample | Sample ID | Sample type | mg used | Final concentration (ng/µl) | Total yield (ng) |
|---|---|---|---|---|---|
| 1 | BenGla-1 | Tail/muscle | 30 | 17.5 | 2 537.5 |
| 2 | BenGla-2 | Tail/muscle | 34 | 27.2 | 3 944 |
| 3 | BenGla-3 | Liver/internal organs | 40 | 136 | 19 720 |
| 4 | BenGla-4 | Hears/brain/skin | 40 | 63.2 | 15 800 |

*Supplementary table 2* Genome data for *Benthosema glaciale*

| | | |
|---|---|---|
| **Project accession data** | | |
| Species | *Benthosema glaciale* | |
| Specimen | fBenGla1 | |
| NCBI taxonomy ID | 125796 | |
| BioProject | XXXXXX | |
| BioSample ID | | |
| Isolate information | | |
| **Raw data accessions** | | |
| PacBio HiFi reads | 3 PACBIO_SMRT (Revio) runs: 3.2 M reads, 34 Gb | |
| Hi-C Illumina reads | 1 ILLUMINA (Illumina NovaSeq S4) run: 426 M pairs of reads, 129 Gb | |
| **Genome assembly metrics** | | |
| HiFi read coverage | 26x | |
| Assembly accession | XXXXXX | |
| Assembly identifier | fBenGla1.1.hap1 | fBenGla1.1.hap2 |
| Span (Mb) | 1282 | 1301 |
| Number of contigs | 6107 | 696 |
| Contig N50 length (Mb) | 0.48 | 0.47 |
| Longest contig (Mb) | 4.4 | 3.3 |
| Number of gaps | 3808 | 3938 |
| Number of scaffolds | 2299 | 3031 |
| Scaffold N50 length (Mb) | 45.0 | 45.0 |
| Longest scaffold (Mb) | 80.9 | 85.5 |

**Commented [MS1]:** Husk senere

**Commented [MS2]:** Husk senere

| | | | |
|---|---|---|---|
| Consensus quality (QV) compared to Hi-C (compared to HiFi) | | 40.17 (51.65) | 39.98 (51.54) |
| Both assemblies | | 40.07 (51.59) | |
| *k*-mer completeness (percentage; compared to HiFi) | | 75.97 (80.87) | 76.54 (81.33) |
| Both assemblies | | 93.85 (98.77) | |
| Percentage of assembly mapped to chromosomes | | 83.00 | 82.58 |
| Comparisons (hap2 aligned to hap1) | Bases in alignment | 372185082 | |
| | Substitutions (percentage) | 1840765 (0.49) | |
| | 1bp deletions | 91033 | |
| | 1bp insertions | 90223 | |
| | 2bp deletions | 54586 | |
| | 2bp insertions | 54553 | |
| | [3,50) deletions | 203842 | |
| | [3,50) insertions | 203669 | |
| | [50,1000) deletions | 26931 | |
| | [50,1000) insertions | 27396 | |
| | >=1000 deletions | 3912 | |
| | >=1000 insertions | 4148 | |
| **Genome annotation metrics** | | | |
| Number of protein-coding genes | | 33856 | 34138 |
| Number of protein-coding genes with functional domain** | | 31972 | 32255 |
| Number of protein-coding genes with gene names | | 23597 | 24085 |

| | | |
|---|---|---|
| BUSCO* | C:95.2%[S:79.8%,D:15.4%],F:1.1%,M:3.7%,n:3640 | C:95.4%[S:79.1%,D:16.3%],F:0.9%,M:3.7%,n:3640 |

* BUSCO scores based on the actinopterygii BUSCO set using v5.7.1. C = complete [S = single copy, D = duplicated], F = fragmented, M = missing, n = number of orthologues in comparison.
**Number of genes annotated with a functional domain as found by InterProScan

*Supplementary table 3* Software tools, versions and sources

| Software tool | Version | Source |
|---|---|---|
| BlobToolKit | 4.1.7 | https://github.com/blobtoolkit/blobtoolkit |
| blobtk | 0.5.8 | https://github.com/blobtoolkit/blobtk |
| BUSCO | 5.4.7 | https://gitlab.com/ezlab/busco |
| hifiasm | 0.19.8 | https://github.com/chhylp123/hifiasm |
| KMC | 3.1.2 | https://github.com/refresh-bio/KMC |
| GenomeScope | 2.0 | https://github.com/tbenavi1/genomescope2.0 |
| HiFiAdapterFilt | 2.0.0 | https://github.com/sheinasim/HiFiAdapterFilt |
| PretextView | 0.2.5 | https://github.com/wtsi-hpag/PretextView |
| PretextMap | 0.1.9 | https://github.com/wtsi-hpag/PretextMap |
| PretextSnapshot | | https://github.com/wtsi-hpag/PretextSnapshot |
| meryl | 1.3.0 | https://github.com/marbl/meryl |
| BWA-MEM | 0.7.17 | https://github.com/lh3/bwa |
| samtools | 1.17 | https://github.com/samtools/samtools |
| YaHS | 1.2a.2 | https://github.com/c-zhou/yahs |
| FCS-GX | 0.4.0 | https://github.com/ncbi/fcs |
| Merqury | 1.3 | https://github.com/marbl/merqury |
| AGAT | 1.0 | https://github.com/NBISweden/AGAT |
| Oatk | 1.0 | https://github.com/c-zhou/oatk |
| miniprot | 0.13 | https://github.com/lh3/miniprot |
| GALBA | 1.0.6 | https://github.com/Gaius-Augustus/GALBA |
| RED | 2018.09.10 | http://toolsmith.ens.utulsa.edu/ |

| | | |
|---|---|---|
| Funannotate | 1.8.17 | https://github.com/nextgenusfs/funannotate |
| EvidenceModeler | 1.1.1 | https://github.com/EVidenceModeler/EVidenceModeler |
| DIAMOND | 2.1.8 | https://github.com/bbuchfink/diamond |
| InterProScan | 5.62-94 | https://www.ebi.ac.uk/interpro/search/sequence/ |
| EMBLmyGFF3 | 2.2 | https://github.com/NBISweden/EMBLmyGFF3 |
| Rapid curation 2.0 | 964d17e997e00c69f25940cf96d3658bda631147 | https://github.com/Nadolina/Rapid-curation-2.0, |

### Supplementary phylogenies in PDF:

Neighbor-joining tree on MHCI Z lineage protein sequences with Poisson substitution model, pairwise deletion and 500 bootstrap replicates made using MEGAX. Myctophiformes species in color. Reference MHCI Z sequences from Grimholt et al 2015.

Neighbor-joining tree on MHCI U lineage protein sequences with Poisson substitution model, pairwise deletion and 500 bootstrap replicates made using MEGAX. Myctophiformes species in color. Reference MHCI U sequences from Grimholt et al 2015.

Neighbor-joining tree on MHCII A lineage protein sequences with Poisson substitution model, pairwise deletion and 500 bootstrap replicates made using MEGAX. Myctophiformes species in color. Reference Salmon MHCII sequences with annotation from Dijkstra et al 2013 in periwinkle blue.

Neighbor-joining tree on MHCII B lineage protein sequences with Poisson substitution model, pairwise deletion and 500 bootstrap replicates made using MEGAX. Myctophiformes species in color. Reference Salmon MHCII sequences with annotation from Dijkstra et al 2013 in periwinkle blue.

### Supplementary data 1

Excel file with all gene findings, scaffold, start, stop.
