## Supplementary_phylogenies for "The genome of the Glacier lanternfish shows loss of MHC I and II function and provides insight into evolution of lanternfish immune systems"

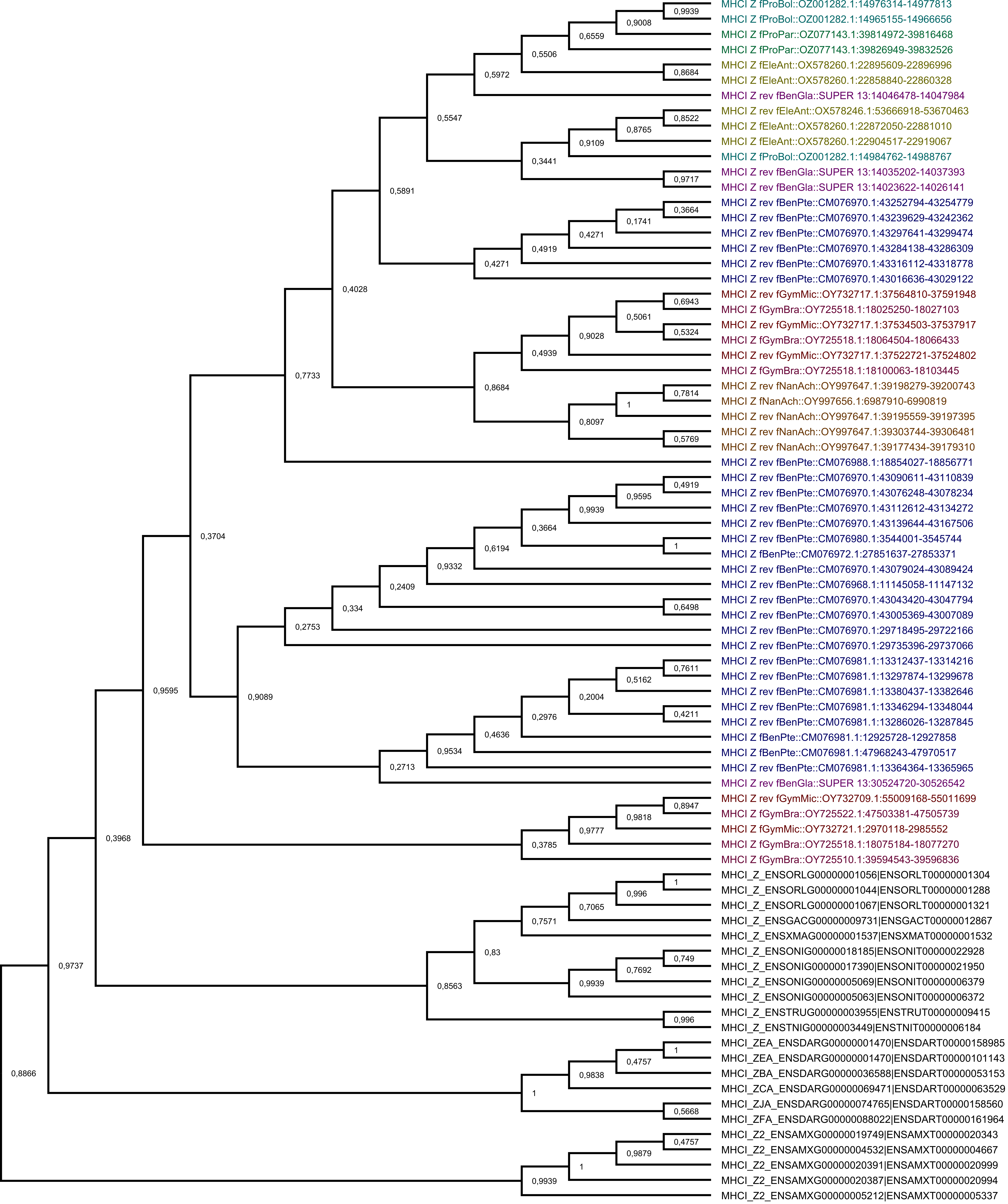


***Supplementary phylogeny:***

***Neighbor-joining tree on MHCI Z lineage protein sequences with Poisson substitution model, pairwise deletion and 500 bootstrap replicates made using MEGAX. Myctophiformes species in color. Reference MHCI Z sequences with annotation from Grimholt et al. 2015.***

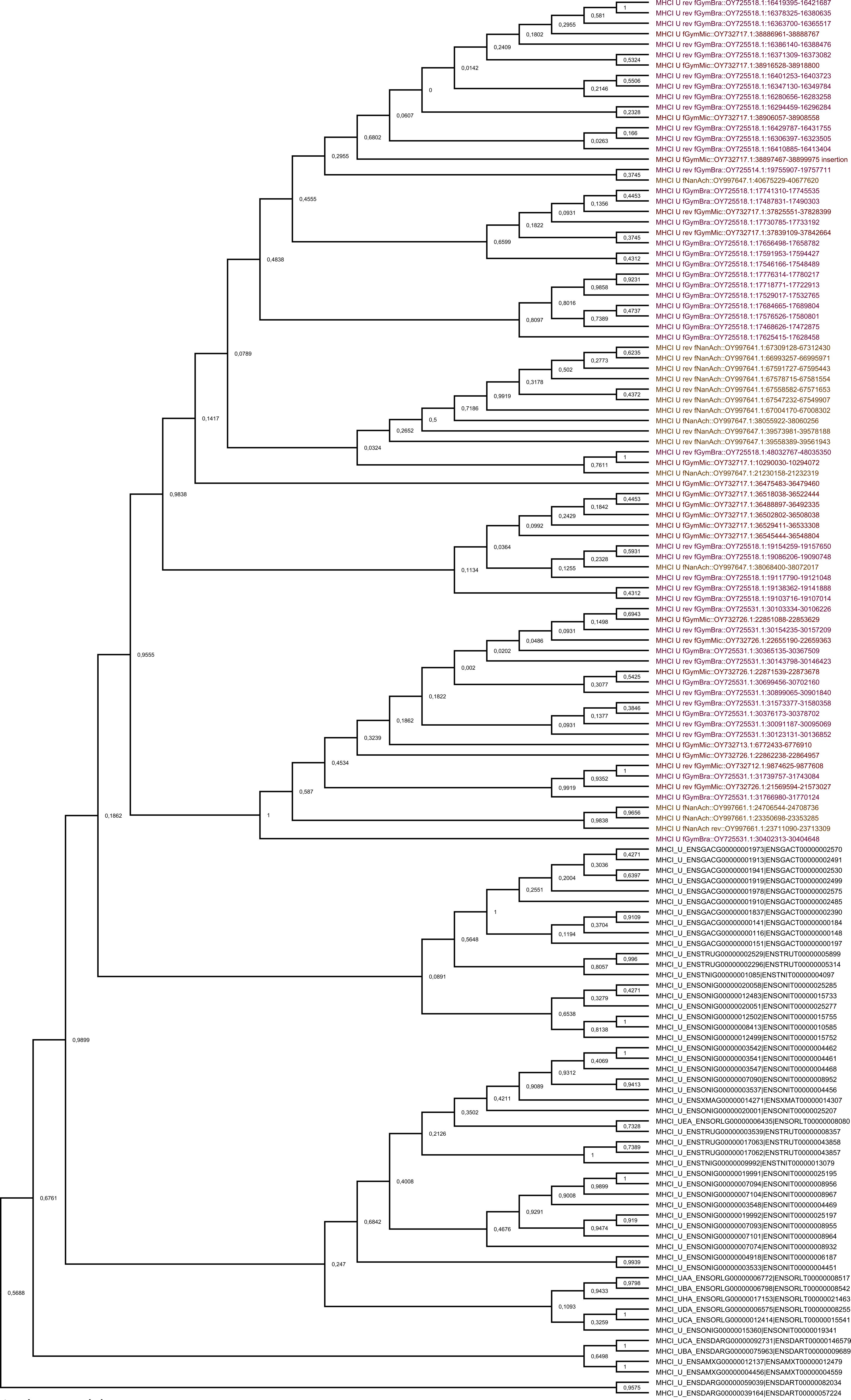


***Supplementary phylogeny:***

***Neighbor-joining tree on MHCI U lineage protein sequences with Poisson substitution model, pairwise deletion and 500 bootstrap replicates made using MEGAX. Myctophiformes species in color. Reference MHCI U sequences with annotation from Grimholt et al. 2015.***

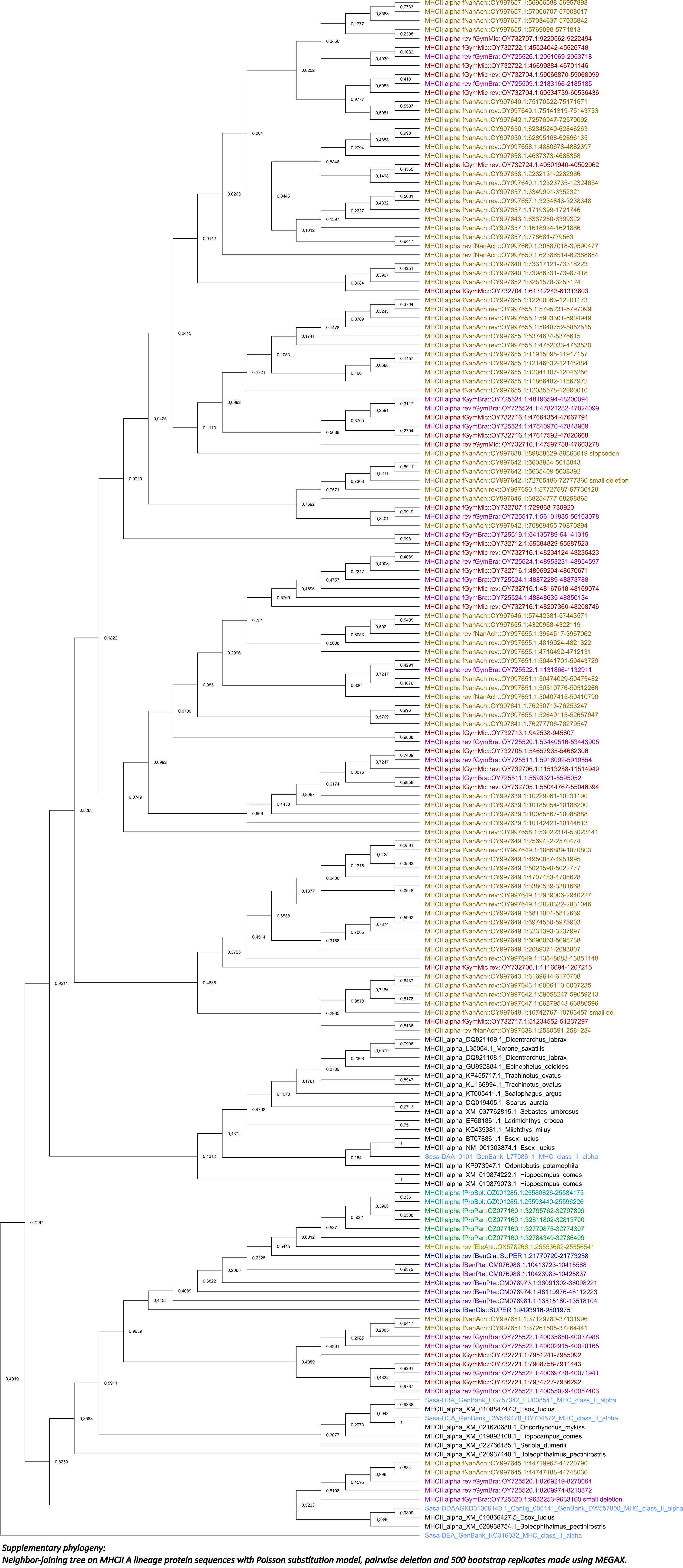


***Supplementary phylogeny:***
***Neighbor-joining tree on MHCII A lineage protein sequences with Poisson substitution model, pairwise deletion and 500 bootstrap replicates made using MEGAX.***
***Myctophiformes species in color. Reference Salmon MHCII sequences with annotation from Dijkstra et al. 2013 in periwinkle blue.***

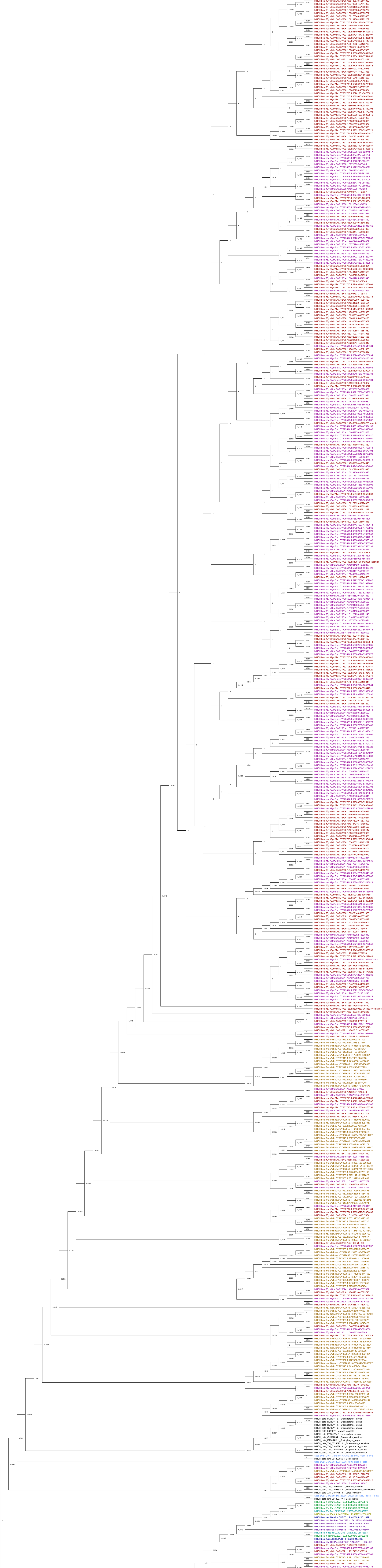

*Supplementary phylogeny:*
*Neighbor-joining tree on MHCII B lineage protein sequences with Poisson substitution model, pairwise deletion and 500 bootstrap replicates made using MEGAX.*
*Myctophiformes species in color. Reference Salmon MHCII sequences with annotation from Dijkstra et al. 2013 in periwinkle blue.*
